# Myeloid hypoxia-inducible factor HIF1A provides cardio-protection during ischemia and reperfusion via induction of netrin-1

**DOI:** 10.1101/2022.04.08.485755

**Authors:** Ka Lin Heck-Swain, Jiwen Li, Wei Ruan, Xiaoyi Yuan, Yanyu Wang, Michael Koeppen, Holger K. Eltzschig

## Abstract

The transcription factor hypoxia-inducible factor HIF1A elicitics cardioprotection from ischemia and reperfusion injury. Here, we investigated tissue-specific pathways that are critical for HIF1A-elicited tissue protection. Initial studies showed that mice with induced global deletion of *Hif1a* (*Hif1a*^*loxP/loxP*^ UbiquitinCre+) have exaggerated myocardial injury during in situ ischemia and reperfusion. Surprisingly, this phenotype was mirrored only in mice with myeloid-specific *Hif1a* deletion *(Hif1a*^*loxP/loxP*^ LysM Cre*+)*. In contrast, mice with myocardial specific (*Hif1a*^*loxP/loxP*^ Myosin Cre*+)*, or vascular *Hif1a* deletion (*Hif1a*^*loxP/loxP*^ VEcadherin Cre+) experienced similar injury levels as controls. Subsequent studies using adoptive transfer of *Hif1a-*deficient polymorphonuclear neutrophils (PMNs) prior to myocardial injury demonstrated increased reperfusion injury. In contrast, adoptive transfer of PMNS treated ex-vivo with the HIF stabilizer dimethyloxalylglycine (DMOG) was associated with attenuated myocardial injury. Moreover, cardioprotection mediated by DMOG was abolished in *Hif1a*^*loxP/loxP*^ LysM Cre+ mice, but not in *Hif2a*^*loxP/loxP*^ LysM Cre+ mice. Finally, studies of PMN-dependent HIF1A target genes implicated the neuronal guidance molecule netrin-1 in mediating the cardioprotective effects of myeloid HIF1A. Taken together, the present studies identified a functional role for myeloid-expressed HIF1A in providing cardio-protection during ischemia and reperfusion injury, which - at least in part - is mediated by the induction of neuronal guidance molecule netrin-1 in neutrophils.

## Introduction

Myocardial infarction remains one of the leading causes of death worldwide. Most commonly, it is an aftereffect of a coronary artery occlusion leading to myocardial ischemia (1). As a consequence of ischemia, myocardial tissue develops profound hypoxia (2), which leads to tissue necrosis if blood flow is not reestablished timely. During myocardial ischemia, tissue hypoxia becomes a strong transcriptional stimulus that activates a coordinated transcriptional program that promotes increased resistance to ischemia and repefusion injury (3, 4). A central role in the coordination of this adaptive response is mediated by stabilization of hypoxia-inducible transcription factor HIF1A (5).

Under normoxic conditions, HIF1A is constantly degraded via the proteasomal pathway due to hydroxylation by prolylhydroxylases (PHDs), causing subsequent binding of the von Hippel Lindau gene product, which targets HIF1A for degradation (6, 7). However, when oxygen levels fall, PHDs become inactivated and HIF1A is rapidly stabilized, translocates to the nucleus and binds to the promoter region of HIF target genes to induce their transcription (2, 3). HIF1A is known to regulate a wide array of genes. For example, gene expression profiles in endothelial cells exposed to normoxia or hypoxia indicate that over 600 target genes are regulated by HIF1A, including gene repression or induction (8). In cardiomyocytes, HIF1A increases the expression of glycolytic pathway enzymes, thereby enhancing their capacity to generate ATP under anaerobic condition and promoting their ability to cope with ischemic tissue injury (9). In myeloid cells, HIF1A transcriptional activity can regulate cell functions, or drive the release of cardio-protective HIF1A target genes (10–12), which can significantly influence inflammatory processes (13). For example, HIF1A stabilization in hypoxia increases the life span of the usually short-lived polymorphonuclear neutrophils (14). Furthermore, HIF1A stabilization increases the motility of myeloid cells towards a chemotactic gradient, bringing them to the site of injury (15, 16).

Based on studies implicating HIF1A stabilization in tissue adaptation and protection during hypoxia or inflammation (3, 17, 18), research efforts were sparked to utilize this pathway as a therapeutic target. Several pharmaceutical companies were successful in finding small molecule inhibitors for PHDs that can be used as HIF activators. In fact, several recent phase-3 clinical trials successfully tested the PHD inhibitors roxadustat or vadadustat for the treatment of renal anemia (19–22). Moreover, experimental studies provide evidences that PHD inhibitors elicit cardioprotection from ischemia and reperfusion injury (5, 23). Together, these studies highlight that the HIF1A pathway could be utilized as a therapeutic target for myocardial ischemia and reperfusion injury.

Cardioprotection by HIF1A is supported by several previous studies (24–26). However, the cellular source of HIF-mediated cardio-protection remains unknown. HIF1A is ubiquitously expressed, including cells involved in myocardial ischemia and reperfusion injury, such as cardiomyocytes, vascular endothelial and myeloid cells (23, 27). To uncover the individual contributions of different cellular compartments during HIF1A-dependent cardio-protection, we used transgenic mice with a *Hif1a* gene flanked by a LoxP site. Breeding these mice with a tissue-specific expression of Cre-recombinase, we stepwise deleted *Hif1a* from different cellular compartments. Surprisingly, we found that the predominant source of HIF1A mediated cardioprotection involves myeloid-cells and uncovered a novel contribution of polymorphonuclear neutrophil (PMNs).

## Materials and Methods

### Mice

All animal procedures were performed in an American Association for the Accreditation of Laboratory Animal Care–accredited facility and approved by the University of Colorado Denver and the University of Texas Health Science Center at Houston (UTHealth) Institutional Animal Care and Use Committee. For all studies, we used mice with an age of 8–16 weeks. C57BL/6J mice were purchased from The Jackson Laboratory (Bar Harbor, ME). *Hif1a*^*loxP/loxP*^ (B6.129-*Hif1a*^*tm3Rsjo*^/J) (28), *Hif2a*^*loxP/loxP*^ (Epas1tm1Mcs/J) (23), *Ntn1*^*loxP/loxP*^ (B6.129(SJL)-Ntn1tm1.1Tek/J) (12), tamoxifen-inducible Ubiquitin Cre+ (B6.Cg-*Ndor1*^*Tg(UBC-cre/ERT2)1Ejb*^/1J) (29), LysM Cre+ (B6.129P2-*Lyz2*^*tm1(cre)Ifo*^/J) (30), VE-cadherin-Cre+ (B6.Cg-Tg(Cdh5-cre)7Mlia/J) (31, 32), and tamoxifen-inducible Myosin-Cre+ (B6.FVB(129)-*A1cf*^*Tg(Myh6-cre/Esr1*)1Jmk*^/J) (32) mice were purchased from The Jackson Laboratory. To obtain tissue-specific mice, we crossbred *Hif1a*^*loxP/loxP*^, *Hif2a*^*loxP/loxP*^ or *Ntn1*^*loxP/loxP*^ mice with tissue-specific Cre-recombinase mouse. For studies using Myosin-Cre+ or Ubiquitin Cre+ mice, mice received tamoxifen dissolved in peanut oil for 5 days (1 mg/day i.p.), followed by a resting period of 7 days prior to experimentation. Mice were genotyped by GeneTyper (New York, NY) or in house per recommended protocol.

### Human neutrophil isolation and cell culture

From healthy subjects, human blood was collected in tubes containing anticoagulant citrate (S-monovette #04.1902, Sarstedt, Nümbrecht, Germany). Neutrophils were isolated using a Percoll-based protocol. Briefly, whole blood was applied to the discontinuous Percoll gradient containing equal vols of 63% and 72% Percoll solution and centrifuged at 400 (x *g*) for 30 min without brake. A mixture of neutrophils and RBCs were collected at the Percoll gradient interface, in which the latter were subsequently removed by isotonic lysis (NH_4_Cl solution). Neutrophils were then pelleted and washed with HBSS(-). Neutrophil purity/viability were assessed by loading a small aliquot into a particle count & size analyzer (Coulter, Backman Coulter) following manufacturer’s instructions. 1-2 × 10^6^ PMN per mL blood were typically isolated with > 95% purity and viability. After isolation, neutrophils were rested 30 min prior to experiments. Cells were cultured 2.5 × 10^6^ cells/mL in RPMI 1640 with 10% FCS at 37 °C in 21% O_2_ (0h timepoint) or 2% O_2_ (hypoxia timepoints).

### Murine model for myocardial ischemia

Detailed procedure of in situ myocardial ischemia and reperfusion injury was previously described (23, 33–35). In brief, we exposed mice to 60 min of myocardial ischemia followed by 2 hours of reperfusion, followed by analysis of infarct size and measurement of cardiac troponin I in the serum. DMOG was given to the mice 4 hours before ischemia started (1mg/mouse, dissolved in normal saline, intraperitoneal). Successful occlusion of the LCA occlusion was confirmed by a color change of the LCA-supplied cardiac tissue. After 60 minutes of ischemia, the weights were removed to allow reperfusion of the tissue. During ischemia, we administered normal saline at a constant rate of 0.1 ml/h. At the beginning of the reperfusion, we increased the infusion rate to 0.6 mL/hour.

### Infarct staining

The detailed procedure to measure the percentage of infarcted area relative to the area undergoing ischemia (area-at-risk) using a counter-stain technique was previously described (36, 37). At the end of the experiment, circulation was flushed with 5 ml of normal saline and LCA was permanently occluded. Following the injection of Evan’s blue dye (600ul, 1%), the heart was excised by the heart basis and kept at −20°C for 15 minutes. Hearts were then sliced and incubated with 1% triphenyltetrazolium chloride (TTC) for 10 minutes at 37°C before fixation in 4% neutral-buffered formalin. To prevent bleaching of the dye, photographs of the heart slices were imaged on the day after the experiments. For this, the tissue slices were placed between glass slides and pictures were taken using a Nikon D5300 camera at 32x magnification. AAR and infarct size was measured by ImageJ software (National institutes of health, USA; Version 1.51k). If an air bubble was unintentionally injected into the cardiac circulation during Evan’s blue injection, resulting in incomplete AAR demarcation, sample was excluded.

### Cardiac troponin I (cTnI) ELISA

As an additional line of evidence for myocardial injury, cTnI was measured by enzyme-linked immunosorbent assay (ELISA), as described (38) using cTnI ELISA Kit (Life Diagnostic, #CTNI-1-HS).

### Neutrophil depletion and adoptive transfer

Untouched PMNs were isolated from bone marrow in donor mice of the indicated genotype (age 6–8 wk) using EasySep mouse neutrophil enrichment kit (Stemcell Technologies Inc.). In a subset of experiment PMNs from C57/J animals were incubated with 1 mM Dimethyloxalylglycine (DMOG; Sigma-Aldrich, #D3695) or vehicle solution for 60 min. Then, 1 × 10^6^ cells were injected into the neutrophil-depleted mice via an arterial catheter over a 10 min period 60 min prior to myocardial ischemia. The detailed procedure for neutrophil depletion in vivo was described previously (39) using Ly6G-specific mAb 1A8 (BioXCell) antibody.

### Immunoblotting experiments

To study the expression of Netrin-1, immunoblotting methods were conducted. At the end of each timepoint, neutrophils in the media were collected and centrifuged at 500 (x *g*) for 5 min to collect the cells. Cells were immediately lysed in IP lysis buffer (Pierce #87788, ThermoFisher) on ice for 30 min then centrifuged at 12,000 (x g) for 10 min. Protein concentration of the lysates was measured by BCA Assay (#23225, ThermoFisher) then incubated in Laemmli sample buffer containing 2-mercaptoethanol at 95 °C for 5 min. Lysate proteins (40 μg) were separated on 7.5 % SDS polyacrylamide gels by SDS-PAGE then transferred onto PVDF membranes using a Bio-Rad Mini Trans-Blot cell. Netrin-1 (LSBio, LS-C743016) and β-actin (Santa Cruz, sc-130656), used as the loading control, were detected using specific primary antibodies. The membranes were subsequently incubated with a secondary antibody coupled to HRP conjugate (Invitrogen #65-6120). The membranes were incubated with ECL reagents (Clarity Max, Biorad and Immunocruz, Santa Cruz, respectively) and the chemiluminescent signal detected by Fusion Imaging System (Vilber Lourmat). Protein expression was quantified by densitometry using ImageJ (National institutes of health, USA; Version 1.51k) and normalized to the expression of β-actin.

### Statistical Analysis

N numbers were pre-determined by power analysis to detect mean difference of 20% with 8% standard deviation (SD). Outliers were detected by Grubb’s test and removed from data. All data approximately followed normal distribution and were ploted as mean±SD. Experimental data were analyzed by unpaired, 2-sided student *t* test or ANOVA with Tukey test for multiple comparison. p value less than 0.05 was considered statistically significant. Statistical analyses were performed by using Prism (version 9.1.2, GraphPad Software Inc.).

## Results

### Global induced deletion of Hif1A is associated with increased myocardial ischemia and reperfusion injury

Previous studies have implicated HIF1A in cardio-protection from ischemia and reperfusion injury (5, 23, 40). For example, mice with heterozygote *Hif1a* deletion are not protected by ischemic preconditioning (41). Similarly, pharmacologic stabilizers of HIF such as DMOG provide robust cardio-protection during ischemia and reperfusion (5). However, the fact that mice with homozygote deletion of *Hif1a* die during embryogenesis (42) have made it challenging to provide direct genetic evidence for *Hif1a* during cardio-protection. Thus, we generated mice with induced global deletion of *Hif1a* upon tamoxifen treatment by crossing *Hif1a*^*loxP/loxP*^ mice with Ubiquitin Cre+ mice to generate *Hif1a*^*loxP/loxP*^ Ubiquitin Cre+ mice (43). To achieve global induced *Hif1a* deletion, mice were treated daily with tamoxifen for 5 days (1 mg i.p./day) and recovered for 7 days before myocardial ischemia and reperfusion injury, using a previously described in situ model (Fig. 1a) (34). Western blot analysis for Hif1a showed Hif1a protein expression in response to Hif1a-stabilizer treatment (DMOG, 1 mg i.p.) in Ubiquitin Cre+ mice. In contrast, Hif1a protein failed to accumulate in the cardiac tissue in *Hif1a*^*loxP/loxP*^ Ubiquitin Cre+ mice following DMOG treatment (Fig. 1b), suggesting successful deletion of Hif1a in these animals.

**Fig. 1.**
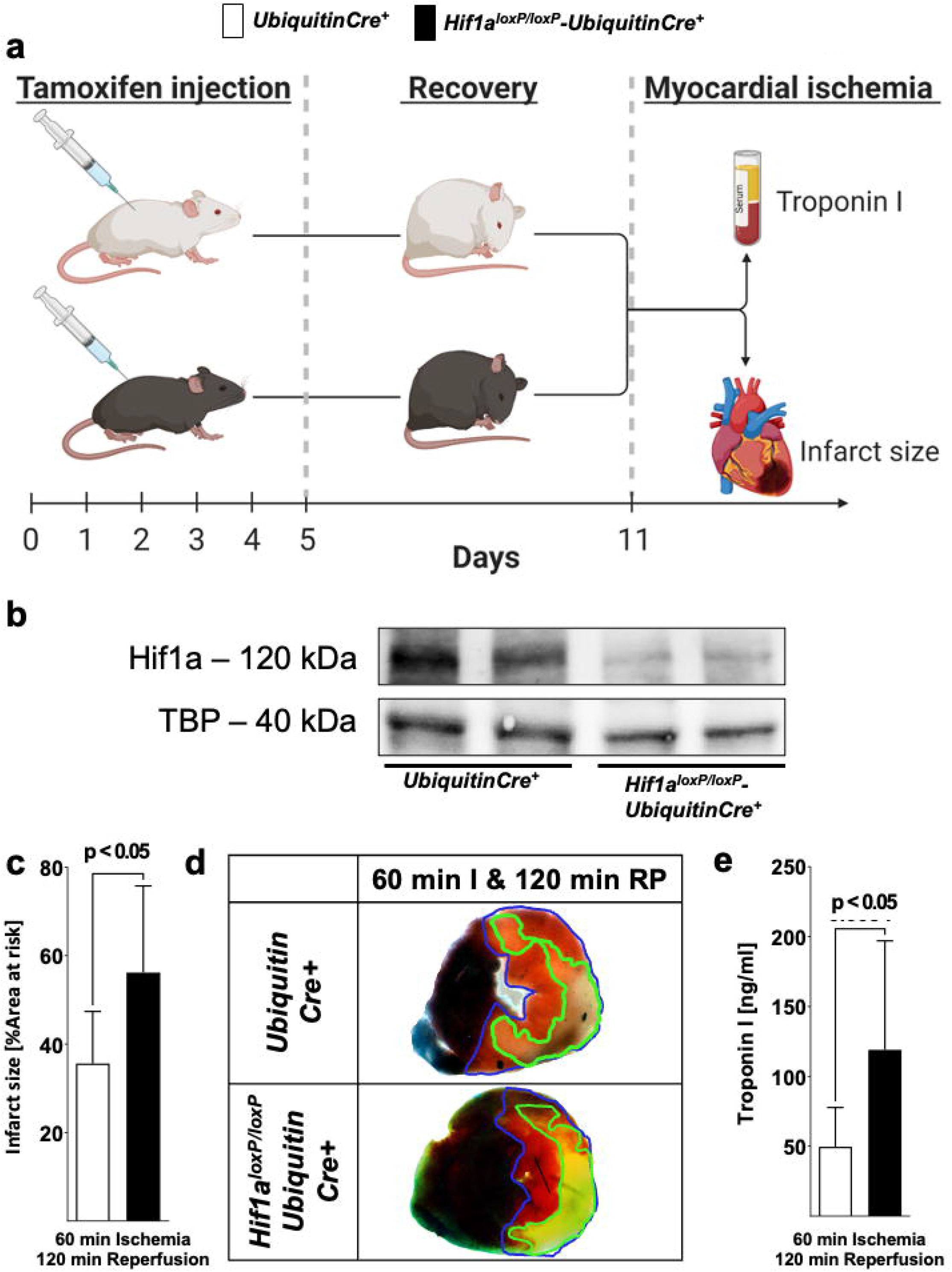
Functional consequences of induced global deletion of *Hif1a* during myocardial ischemia and reperfusion injury. **a** Experimental approach to study global induced *Hif1a*-deficiency. *Hif1a*^*loxP/loxP*^ mice were crossed with Cre-recombinase expressing mice under the control of Ubiquitin promoter (UbiquitinCre+); these mice express cre-recombinase under the control of a tamoxifen-inducer. Gender and weight matched control animals (UbiquitinCre+) or *Hif1a*^*loxP/loxP*^ UbiquitinCre+ mice received a daily dose of 1 mg i.p. tamoxifen 5 consecutive days to induce Cre-recombinase activity. After 7 days, animals underwent experimental protocols (60 min of *in situ* myocardial ischemia followed by 120 min of reperfusion), followed by serum collection to determine concentrations of troponin I and TCC staining for the infarct sizes. **b** Hif1A immunoblot performed in protein isolated from myocardial tissue after treatment with the pharmacologic HIF activator dimethyloxalylglycine (1 mg DMOG i.p.) 4 h prior to tissue collection from UbiquitinCre+ or *Hif1a*^*loxP/loxP*^ UbiquitinCre+. A representative image of 3 individual experiments is presented. **c** Infarct sizes of UbiquitinCre+ or *Hif1a*^*loxP/loxP*^ UbiquitinCre+ mice are displayed as percentage of the area-at-risk after 60 min of ischemia, followed by 120 min of reperfusion (UbiquitinCre+ n=7; *Hif1a*^*loxP/loxP*^ UbiquitinCre+ n=5, per group mean ± SD; statistical analysis by two-sided, unpaireds Student’s t-test; mice were matched by age, gender and weight). **d** Representative infarct staining of UbiquitinCre+ or *Hif1a*^*loxP/loxP*^ UbiquitinCre+ mice (60 min ischemia and 120 min reperfusion). **e** Troponin serum levels after 60 min ischemia, followed by 120 min of reperfusion in UbiquitinCre+ or *Hif1a*^*loxP/loxP*^ UbiquitinCre+ mice (UbiquitinCre+ n=12; *Hif1a*^*loxP/loxP*^ UbiquitinCre+ n=8 presented as mean ± SD; compared by Student’s t-test).

To assess the functional consequences of global induced Hif1a deletion on myocardial ischemia and reperfusion injury, we exposed *Hif1a*^*loxP/loxP*^ Ubiquitin Cre+ mice to in situ myocardial ischemia and reperfusion following tamoxifen treatment and recovery (Fig. 1a) and assessed myocardial injury by infarct sizes and serum troponin levels. These studies demonstrated significantly larger myocardial infarct sizes following 60 min of ischemia and 2h or reperfusion in *Hif1a*^*loxP/loxP*^ Ubiquitin-Cre+ mice as compared to littermate control Ubiquitin Cre+ mice following tamoxifen treatment (Fig. 1c, d). Similarly, *Hif1a*^*loxP/loxP*^ Ubiquitin Cre+ mice experienced significantly elevated of the cardiac injury marker troponin I (Fig. 1e). Together these studies demonstrate that induced deletion of Hif1a is associated with dramatic increases in myocardial injury following ischemia and reperfusion, and confirm previous studies implicating HIF1A in cardioprotection (5, 40).

### Tissue-specific *Hif1a*-deletion reveals a selective role for myeloid-derived Hif1A in mediating cardioprotection during ischemia and reperfusion

Based on the above studies showing the global induced deletion of Hif1A has deleterious effects during in situ myocardial ischemia and reperfusion injury, we next pursued studies to identify tissue-specific contributions of *Hif1a* in cardioprotection. For this purpose, we generated mice with *Hif1a* deletion in different tissue compartments, including cardiomyocytes (*Hif1a*^*loxP/loxP*^ Myosin Cre+ mice) (23), vascular endothelial cells (*Hif1a*^*loxP/loxP*^ VEcadherin Cre+ mice) (27) or myeloid cells (*Hif1a*^*loxP/loxP*^ LysM Cre+ mice) (44). We subjected these mouse lines with respective gender, weight and age matched Cre+ control mice to 60 min of myocardial ischemia and 120 min of reperfusion and measured myocardial injury by assessing infarct size area and serum troponin levels, as we have done previously (12, 23, 35, 45). Consistent with previous studies (23), we found that *Hif1a*^*loxP/loxP*^ Myosin Cre+ mice had similar infarct sizes and troponin I levels compared to Myosin Cre+ controls (Fig. 2a-c). Similarly, deletion of Hif1a in vascular endothelia (*Hif1a*^*loxP/loxP*^ VEcadherin Cre+ mice) did not alter the susceptibility to myocardial injury (Fig. 2d-f). Surprisingly, we found the predominant phenotype in mice with deletion of *Hif1a* in the myeloid compartment. Indeed, *Hif1a*^*loxP/loxP*^ LysM Cre+ mice had dramatic increases in myocardial infarct sizes, and increased troponin I leakage into the plasma, as compared to LysM Cre+ control animals (Fig. 2g-i). Together, these studies provide novel evidence that mice with myeloid-specific *Hif1a* deletion experience increased susceptibility to myocardial ischemia and reperfusion injury, essentially resembling the findings in mice with induced global deletion of Hif1A (Fig. 1c-e).

**Fig. 2.**
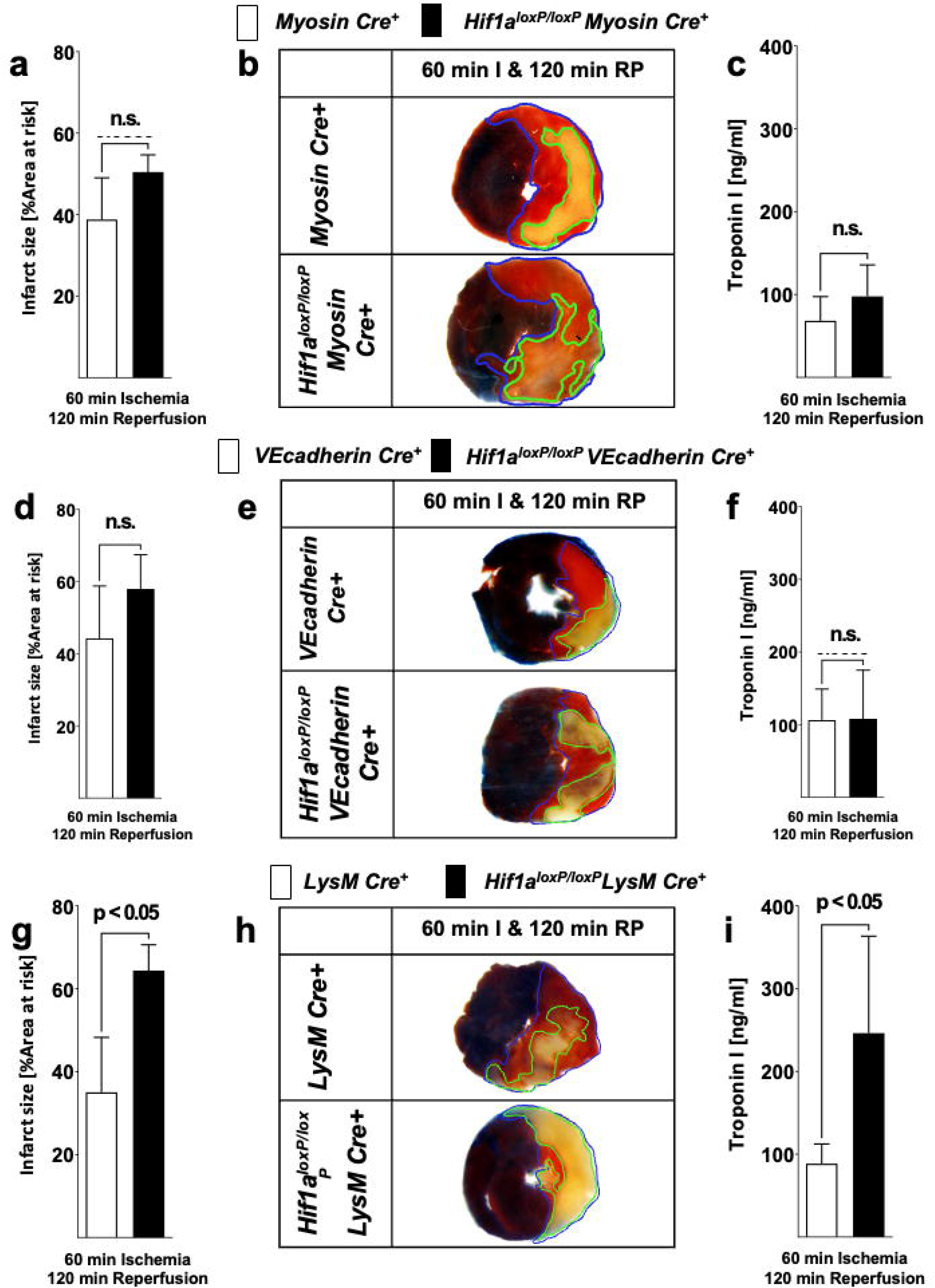
Myocardial ischemia and reperfusion injury in mice with tissue-specific Hif1a deletion in cardiac myocytes, vascular endothelia or myeloid inflammatory cells. **a-i** *Hif1a*^*loxP/loxP*^ were crossed with mice expressing Cre-recombinase under the control of heavy chain myosin promoter including a tamoxifen inducer (*Hif1a*^*loxP/loxP*^ Myosin Cre+ mice) (23), vascular endothelial cadherin (*Hif1a*^*loxP/loxP*^ VEcadherin Cre+ mice) (27). (*Hif1a*^*loxP/loxP*^ VEcadherin Cre^+^) or lysozyme 2 (*Hif1a*^*loxP/loxP*^ LysM Cre+ mice) (44). Mice underwent 60 min myocardial ischemia, followed 120 min reperfusion. Myocardial injury was determined by area-at-risk and troponin I serum concentration **a** *Hif1a*^*loxP/loxP*^ Myosin Cre+ mice and age, gender and weighed matched Myosin Cre+ mice received 1 mg i.p. tamoxifen per day over 5 days followed by 7 days of recovery prior to experimentation. Infarct sizes ± SD in Myosin Cre+ or *Hif1a*^*loxP/loxP*^ Myosin Cre*+* presented as percentage to the area-at-risk after 60 min of ischemia, followed by 120 min of reperfusion (Myosin Cre+ n=6; *Hif1a*^*loxP/loxP*^ Myosin Cre+ n=4). **b** Representative infarct staining from Myosin Cre+ or *Hif1a*^*loxP/loxP*^ Myosin Cre+ (60 min ischemia and 120 min reperfusion). **c** Troponin serum levels after 60 min ischemia, followed by 120 min of reperfusion in Myosin Cre+ or *Hif1a*^*loxP/loxP*^ Myosin Cre+ (Myosin Cre+ n=5; *Hif1a*^*loxP/loxP*^ Myosin Cre+ n=5). **d** Infarct sizes ± SD in VEcadherin Cre+ or *Hif1a*^*loxP/loxP*^ VEcadherin Cre+ presented as percentage to the area-at-risk after 60 min of ischemia, followed by 120 min of reperfusion (VEcadherin Cre+ n=7; *Hif1a*^*loxP/loxP*^ VEcadherin Cre+ n=7). **e** Representative infarct staining from VEcadherin Cre+ or *Hif1a*^*loxP/loxP*^ VEcadherin Cre+ (60 min ischemia and 120 min reperfusion). **f** Troponin serum levels after 60 min ischemia, followed by 120 min of reperfusion in VEcadherin Cre+ or *Hif1a*^*loxP/loxP*^ VEcadherin Cre+ (VEcadherin Cre^+^ n=4; *Hif1a*^*loxP/loxP*^ VEcadherin Cre+ n=6). **g** Infarct sizes ± SD in LysM Cre+ or *Hif1a*^*loxP/loxP*^ LysM Cre+ presented as percentage to the area-at-risk after 60 min of ischemia, followed by 120 min of reperfusion (LysM Cre+ n=8; *Hif1a*^*loxP/loxP*^ LysM Cre+ n=4). **h** Representative infarct staining from LysM Cre+ or *Hif1a*^*loxP/loxP*^ LysM Cre+ (60 min ischemia and 120 min reperfusion). **i** Troponin serum levels after 60 min ischemia, followed by 120 min of reperfusion in LysM Cre+ or *Hif1a*^*loxP/loxP*^ LysM Cre+ (LysM Cre+ n=4; *Hif1a*^*loxP/loxP*^ LysM Cre+ n=3); (all values represented as mean ± SD; statistical significance assessed by two-sided, unpaired Student’s t-test).

### Transfer of *Hif1a*-deficient polymorphonuclear neutrophils (PMNs) is associated with more severe myocardial ischemia and reperfusion injury

The above studies indicate myeloid-dependent HIF1A in cardio-protection from ischemia and reperfusion injury. As next step, we pursued studies to further characterize specific cellular populations that are critical in this response. Due to the fact that PMNs are the predominant bone marrow-derived inflammatory cells in the post-ischemic myocardium (38), we next performed studies to address PMN-dependent HIF1A. For this purpose, we performed an adoptive transfer as described previously (45). In short, C57BL6J animals received 250 μg of 1A8 Ly6G-antibody 24 h prior to experimentation to deplete PMNs. On the day of the experiment, we isolated a pure population of mature PMNs from *Hif1a*^*loxP/loxP*^ LysM Cre+ or matched wildtype control mice and infused via an intravascular catheter into PMN-depleted animals followed by 60 min of myocardial ischemia followed by 2h of reperfusion (Fig. 3a). In line with a functional role of HIF1A expressed in PMNs, we found that reconstitution with *Hif1a*-deficient PMNs was associated with dramatically increased infarcts sizes, when compared to mice reconstituted with control PMNs (Fig. 3b-c), resembling myocardial injury phenotypes in *Hif1a*^*loxP/loxP*^ LysM Cre+ mice. Together, these results indicate that PMN-dependent Hif1a deficiency is associated with elevated myocardial ischemia and reperfusion injury and implicates PMN-dependent HIF1A in cardioprotection.

**Fig. 3.**
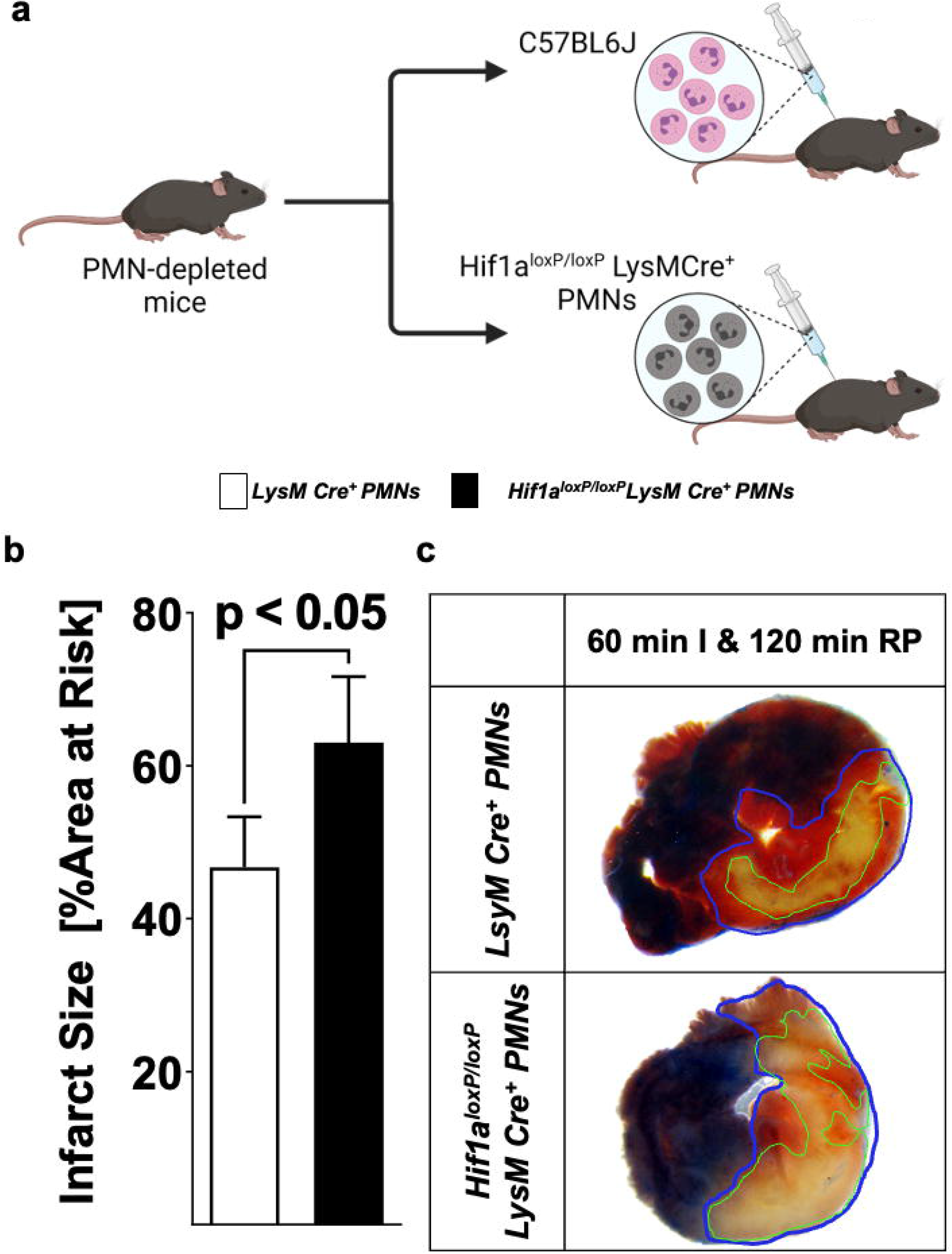
Effects of adoptive transfer of polymorphonuclear neutrophils (PMNs) from *Hif1a*^*loxP/loxP*^ LysM Cre+ into wildtype mice during myocardial ischemia and reperfusion injury. **a** Schematic of adoptive transfer model. C57BL6J or Hif1a-deficient PMNs (*Hif1a*^*loxP/loxP*^ LysM Cre+) were isolated by negative selection and transferred into PMN-depleted C57BL6 animals (1A8 Ly6G-sepcific antibody 24 hours prior to experiment). Then animals underwent 60 min of ischemia, followed by 120 min of reperfusion with assessment of myocardial injury. **b** Infarct sizes ± SD in mice receiving PMNs from C57BL6J or *Hif1a*^*loxP/loxP*^ LysM Cre+ mice presented as percentage to the area-at-risk after 60 min of ischemia, followed by 120 min of reperfusion (LysM Cre+ PMNs n=4; *Hif1a*^*loxP/loxP*^ LysM Cre+ n=4; all values represented as mean ± SD; statistical significance assessed by two-sided, unpaired Student’s t-test). **c** Representative infarct staining from neutrophil-depleted mice receiving PMNs from LysM Cre+ or *Hif1a*^*loxP/loxP*^ LysM Cre+ mice.

### Pharmacologic stabilization of HIF in isolated PMNs prior to myocardial injury confers cardioprotection

Based on the above studies showing that reconstitution with Hif1A-deficient PMNs prior to myocardial injury is associated with more severe myocardial infarction after ischemia and reperfusion, we next pursued opposite studies using PMNs treated ex vivo with a pharmacologic HIF activator. For this purpse, we used the HIF stabilizer DMOG. Previous studies from our laboratory had shown that systemic treatment with DMOG provides cardioprotection via stabilization of HIF1A (5, 23, 40). Similar to the experimental approach above, we first pursued PMN depletion prior to the experimentation by injecting mice with 250 μg of 1A8 Ly6G-antibody 24 h prior to myocardial ischemia and reperfusion. We isolated PMNs from C57BL6J mice and subsequently exposed PMNs for 60 min to vehicle or 1 mM DMOG (46). After the incubation period, we performed an adoptive transfer into PMN-depleted mice followed by 60 min of in situ ischemia and 120 min of reperfusion (Fig. 4a). Mice that had received PMNs pre-treated with DMOG had significantly attenuated myocardial injury as compared to vehicle controls (Fig. 4b-c). Taken together, these studies highlight that adoptive transfer of ex vivo DMOG treated PMNs before myocardial ischemia and reperfusion is associated with attenuated reperfusion injury and suggests a functional role of PMN-dependent HIF in cardioprotection.

**Fig. 4.**
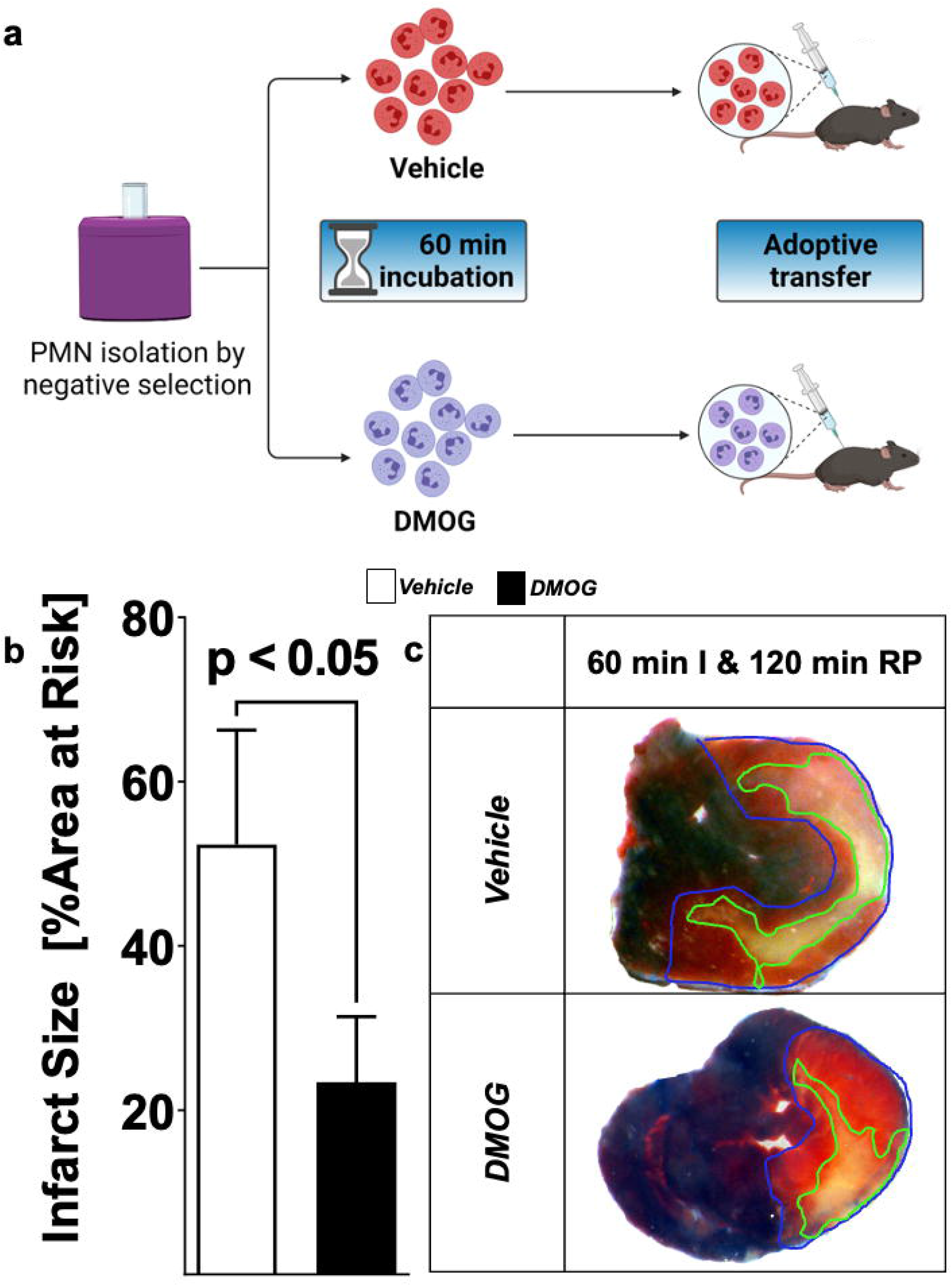
Effect of HIF stabilization in PMNs on myocardial injury. **(a)** Schematic of pharmacological HIF stabilization in PMNs. Cells were harvested by negative selection from C57BL6 mice; isolated cells were incubated for 60 min in vehicle or 1 mM DMOG solution. Then cells were adoptively transferred into PMN-depleted C57BL6, treated with 1A8 Ly6G-sepcific antibody 24 hours prior to experiment. **b** Infarct sizes ± SD in mice receiving vehicle or DMOG treated PMNs presented as percentage to the area-at-risk after 60 min of ischemia, followed by 120 min of reperfusion (Vehicle n=5; DMOG n=4; all values represented as mean ± SD; statistical significance assessed by two-sided, unpaired Student’s t-test). **c** Representative infarct staining from neutrophil-depleted mice receiving PMNs treated with DMOG or vehicle.

### Selective role of myeloid HIF1A deficiency in mediating the cardioprotective effects of PHD-inhibitor treatment DMOG

Previous studies have implicated both isoforms of HIFA (HIF1A or HIF2A) in cardio-protection from ischemia and reperfusion injury, including protective effects mediated by treatment with HIF stabilizers such as DMOG (5, 23, 35). To gain insight into the relative contributions of myeloid-expressed *Hif1a* or *Hif2a* in mediating DMOG elicited cardioprotection, we compared *Hif1a*^*loxP/loxP*^ LysM Cre+ or *Hif2a*^*loxP/loxP*^ LysM Cre+ on their capacity for infarct size reduction provided by pre-treatment with DMOG. In these studies, *Hif1a*^*loxP/loxP*^ LysM Cre+, or *Hif1a*^*loxP/loxP*^ LysM Cre+ were pre-treated with 1 mg DMOG i.p. or vehicle control 4h before myocardial ischemia. Subsequently, animals were exposed to 60 min of myocardial ischemia and 120 min of reperfusion. In line with a functional role of myeloid-expressed Hif1a in mediating DMOG-protection we found that *Hif2a*^*loxP/loxP*^ LysM Cre+ mice are responsive to DMOG pre-treatment with reduced ischemia and reperfusion injury compared to vehicle, as demonstrated in myocardial infarct sizes and troponin I-leakage (Fig. 5a-c). In contrast, we found that *Hif1a*^*loxP/loxP*^ LysM Cre+ failed to respond to pre-treatment with DMOG. Myocardial infarct sized of *Hif1a*^*loxP/loxP*^ LysM Cre+ pre-treated with DMOG were similar to that of *Hif1a*^*loxP/loxP*^ LysM Cre+ pretreated with vehicle control (Fig. 5d-f). Taken together, these results indicate that myeloid HIF1A as opposed to myeloid HIF2A is required for the cardio-protection provided by PHD inhibitors. In conjunction with our studies showing attenuated infarct sizes after ex vivo treatment with HIF stabilizer, these findings provide additional data implicating PMN-dependent HIF1A in cardio-protection from ischemia and reperfusion injury.

**Fig. 5.**
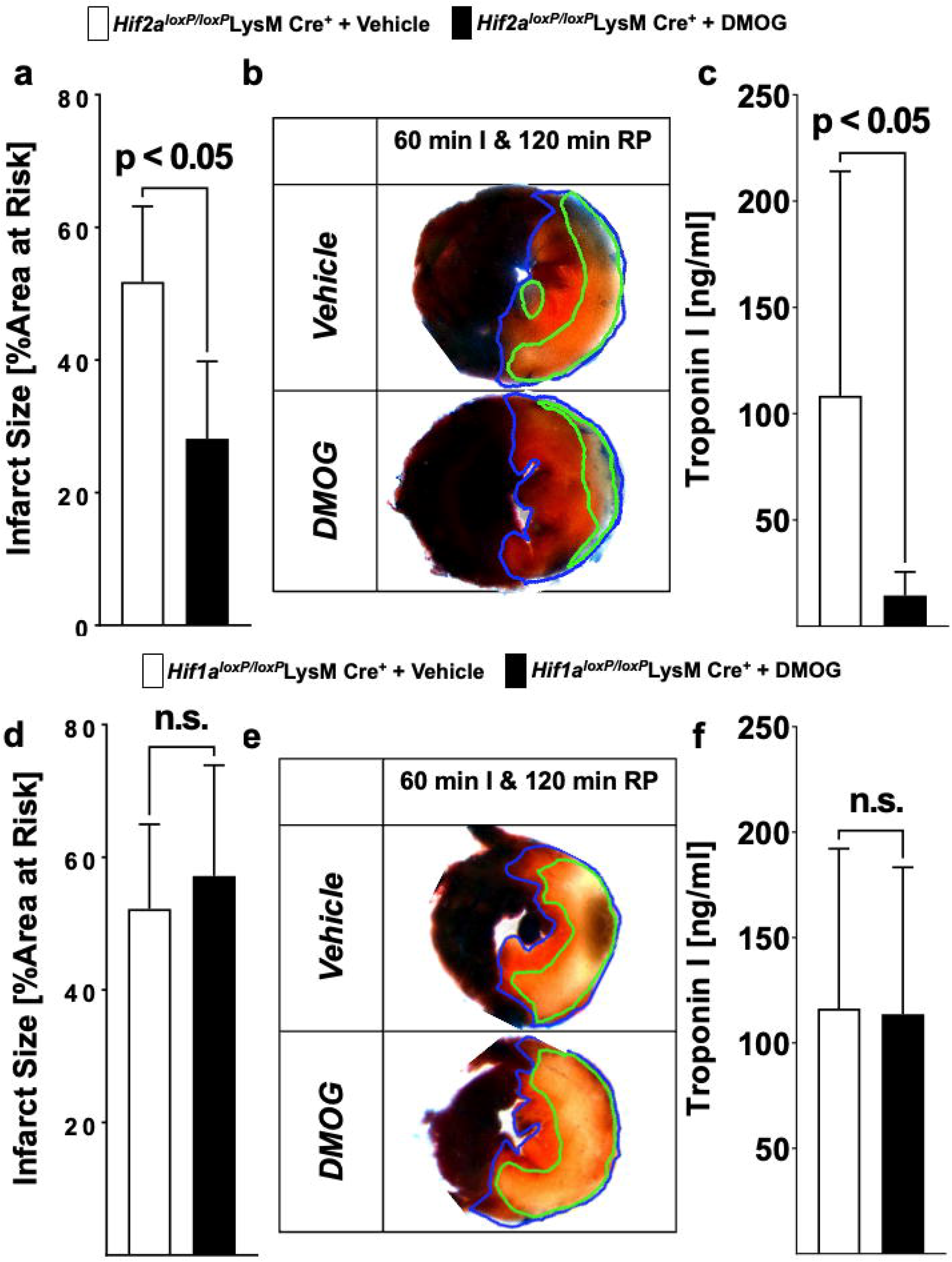
Selective HIF-Isoform deletion in PMNs in myocardial ischemia and reperfusion. **a-f** *Hif1a*^*loxP/loxP*^ or *Hif2a*^*loxP/loxP*^ were crossed with Cre-recombinase expressing mice under the control of the lysozyme 2 promoter (LysM Cre+) to generate myeloid-specific Hif1a- or Hif2a-deficient animals. Mice received vehicle or 1 mg of HIF-stabilizing DMOG 4 hours prior to the ischemia. Mice underwent 60 min of ischemia and 120 min reperfusion and assessment of myocardial injury infarct sizes measurement and serum concentrations of troponin I. **a** Infarct sizes were determined in vehicle or DMOG treated *Hif2a*^*loxP/loxP*^ LysM Cre+ mice (n=5 per group). **b** Representative infarct staining from vehicle or DMOG treated *Hif2a*^*loxP/loxP*^ LysM Cre+ mice. **c** Troponin serum levels in vehicle or DMOG treated *Hif2a*^*loxP/loxP*^ LysM Cre+ mice. (n=5 per group). **d** Infarct sizes were determined in vehicle or DMOG treated *Hif1a*^*loxP/loxP*^ LysM Cre+ mice (*Hif1a*^*loxP/loxP*^LysM Cre+ + Vehicle n=5; *Hif1a*^*loxP/loxP*^LysM Cre+ + DMOG n=6). **e** Representative infarct staining from vehicle or DMOG treated *Hif1a*^*loxP/loxP*^ LysM Cre+ mice. **f** Troponin serum levels in vehicle or DMOG treated *Hif1a*^*loxP/loxP*^ LysM Cre+ mice (*Hif1a*^*loxP/loxP*^LysM Cre+ + Vehicle n=5; *Hif1a*^*loxP/loxP*^LysM Cre+ + DMOG n=6). (All data presented as mean ± SD. Statistical significance assessed by two-sided, unpaired Student’s t-test).

### Identification of neuronal guidance molecule netrin-1 as PMN-dependent Hif1A target critical for HIF-dependent cardioprotection

Based on the above studies demonstrating functional roles of neutrophil-dependent HIF1A in cardioprotection from ischemia and reperfusion, we subsequently pursued studies to identify a transcriptional target in PMNs that would account for these observations. Recent studies from our laboratory had shown that PMN-derived netrin-1 functions to attenuate in situ myocardial ischemia and reperfusion injury. Netrin-1 was originally described as a neuronal guidance molecule important in brain development (47). However, more recent studies implicate netrin in orchestrating inflammatory events, including myocardial inflammation (48). Moreover, other studies had implicated HIF1A in the transcriptional regulation of netrin-1 during hypoxia, and identified netrin-1 as a classic HIF-target gene (44, 49). To examine a functional role of hypoxia-signaling in inducing PMN-dependent netrin-1, we first examined expression of netrin-1 during condition of ambient hypoxia. For this purpose, we freshly isolated human PMNs from peripheral blood and submitted the cells to 0, 2, and 4 h of ambient hypoxia (1% oxygen) and probed the cell lysates for netrin-1 expression by Western Blot. Consistent with previous studies showing induction of netrin-1 expression during hypoxia (49), we found a very robust induction of PMN-dependent netrin-1 during ambient hypoxia (Fig. 6a and b). Recent studies from our laboratory demonstrated the mice with deletion of myeloid-expressed netrin-1 (*Ntn1*^*loxP/loxP*^ LysM Cre+ mice) experience increased myocardial infarct sizes and show that elevated levels of netrin-1 during myocardial infarction predominantly stems from PMNs (12). To further establish a link between the HIF-pathway and myeloid-dependent netrin-1 during cardioprotection, we next pursued studies with the pharmacologic HIF stabilizer DMOG. Here, we hypothesized that if myeloid-dependent HIF functions through induction of netrin-1 to provide cardioprotection, the cardioprotective effects of DMOG would be abolished in *Ntn1*^*loxP/loxP*^ LysM Cre+ mice. To address this hypothesis, we treated *Ntn1*^*loxP/loxP*^ LysM Cre+ mice with DMOG (1mg i.p. 4h prior to myocardial ischemia) or vehicle control, a treatment protocol that we had previously shown to be effective in reducing myocardial infarct sizes (5, 23). Consistent with a functional role of PMN-dependent netrin-1 in mediating DMOG-dependent cardioprotection, we found that the previously shown reduction of myocardial infarct sizes associated with DMOG treatment (Fig. 5a-c) (5, 23) were completely abolished in *Ntn1*^*loxP/loxP*^ LysM Cre+ mice (Fig. 6c-e). Together with recent studies from our laboratory showing that *Ntn1*^*loxP/loxP*^ LysM Cre+ mice experience increased infarct sizes (12), these findings implicate HIF-dependent induction of netrin-1 in the the cardioprotection provided by myeloid-expressed HIF1A during ischemia and reperfusion injury of the heart.

**Fig. 6.**
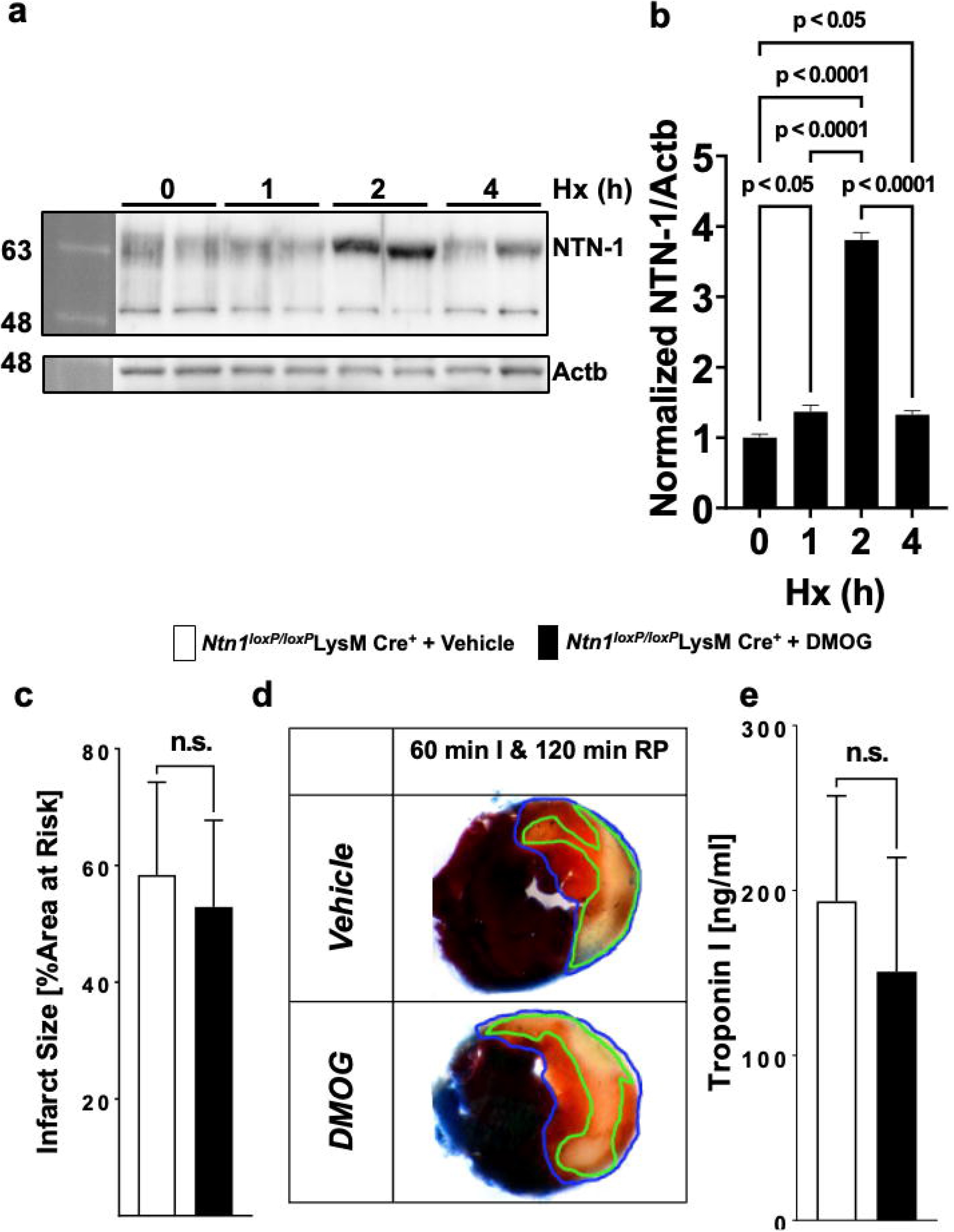
Netrin-1 as PMN-dependent Hif1A target critical for HIF-dependent cardioprotection. **a-b** Isolated human PMNs from healthy donors underwent hypoxia for 0, 2 and 4 hours followed by cell lysis and total protein harvest. **a** Immunoblot for NTN1. b-Actin (Actb) served as loading control. One representative blot out of three experiments is shown. **b** Quantitation by densitometry of the NTN1 immunoblot results relative to b-Actin (n=2 per group; per group mean□±□ SD; compared by one-way ANOVA followed by Tukey’s multiple comparisons test). **c-e** *Ntn-1*^*loxP/loxP*^ *mice* were crossed with Cre-recombinase expressing mice under the control of the lysozyme 2 promoter (LysM Cre+) to generate myeloid-specific Ntn-1-deficient mice. Mice received vehicle or 1 mg DMOG 4 hours prior to the ischemia, then underwent 60 min of ischemia followed by 120 min reperfusion and assessment of myocardial injury. **c** Infarct sizes were determined in vehicle or DMOG-treated *Ntn-1*^*loxP/loxP*^ LysM Cre+ (*Ntn-1*^*loxP/loxP*^ LysM Cre+ with Vehicle n=5; *Ntn-1*^*loxP/loxP*^ LysM Cre+ with DMOG n=6). **d** Representative infarct staining from vehicle or DMOG treated *Ntn-1*^*loxP/loxP*^ LysM Cre+. **e** Troponin serum levels of vehicle or DMOG treated *Ntn-1*^*loxP/loxP*^ LysM Cre+ mice. (*Ntn-1*^*loxP/loxP*^ LysM Cre+ with Vehicle n=5; *Ntn-1*^*loxP/loxP*^ LysM Cre+ with DMOG n=6). All data presented as mean ± SD. Statistical significance assessed by two-sided, unpaired Student’s t-test. *Note: DMOG-induced cardioprotection was absent in Ntn-1*^*loxP/loxP*^ LysM Cre+.

## Discussion

Hypoxia-inducible factors such as HIF1A have been implicated in tissue adaption during limited oxygen availability, inflammation, or ischemia and reperfusion injury (50). While previous studies have directly implicated HIF1a in limiting myocardial infarct sizes during ischemia and reperfusion injury (5, 40, 41, 51, 52), the relative contributions of HIF1A expressed in different cell types in the heart remain elusive. To investigate tissue-specific functions of HIF1A in cardioprotection from in situ ischemia and reperfusion injury, we took a stepwise approach. In initial studies, we used a genetic approach, where Hif1A deletion was induced in the adult mouse, thereby circumventing embryonic lethality of homozygote Hif1a deletion (42). These studies revealed dramatic increase of myocardial infarct sizes in mice with induced global Hif1A deletion (*Hif1a*^*loxP/loxP*^ UbiquitinCre+ mice) as compared to Cre+ controls. As second step, we pursued studies of myocardial ischemia and reperfusion injury in mice with genetic deletion of *Hif1a* in different tissue compartments, including cardiomyocytes (*Hif1a*^*loxP/loxP*^ Myosin Cre*+* mice), vascular endothelial cells (*Hif1a*^*loxP/loxP*^ VEcadherin Cre+ mice) or in myeloid inflammatory cells (*Hif1a*^*loxP/loxP*^ LysM Cre*+* mice). To our surprise, only mice with deletion of *Hif1a* in myeloid inflammatory cells resembled the previous findings we had established in mice with induced global deletion of Hif1A, thereby indirectly implicating myeloid-dependent *Hif1a* in cardio-protection from ischemia and reperfusion. Due to the critical functions and high presence of PMNs during myocardial reperfusion injury (53), we studied mice with reconstitution of wild-type or *Hif1a*-deficient neutrophils following antibody-mediate PMN depletion (54). While mice with *Hif1a*-deficient PMNs experience increased myocardial injury, mice with reconstitution of wild-type PMN that received ex vivo treatment with HIF stabilizer DMOG were protected. Finally, studies that focused on transcriptional targets of HIF1A in neutrophils implicated PMN-dependent netrin-1 in mediating HIF1A elicited cardioprotection. Together, these studies highlight a functional role of myeloid-dependent HIF1A in attenuating myocardial ischemia and reperfusion, and its transcriptional target netrin-1. Based on these studies, using HIF stabilizers or treatment with recombinant netrin-1 may represent novel approaches to treat or prevent myocardial injury during myocardial ischemia and reperfusion.

While the current studies directly implicate HIF1A in attenuating myocardial ischemia and reperfusion injury, other studies have also implicated HIF2A in cardioprotection. HIF2A is an isoform of HIF1A and several studies have identified either complementary or opposing functions to HIF1A (35, 55–58). As such, it is not surprising that previous studies have also identified a functional role of Hif2A in attenuating myocardial infarct sizes during ischemia and reperfusion injury (23). In contrast to the current studies that implicated myeloid-dependent Hif1A in cardioprotection, those studies demonstrated a functional role of HIF2A expressed in cardiac myocytes. In fact, *Hif2a*^*loxP/loxP*^ Myosin Cre+ mice exhibited larger infarct sizes during in situ myocardial ischemia and reperfusion injury (23). Subsequent studies on myocyte-dependent HIF2A target genes implicate the epidermal growth factor amphiregulin (AREG) in mediating HIF2A dependent cardioprotection (23). Additional studies in myocardial biopsies of patients with ongoing ischemia revealed elevated levels of AREG (23), of the amphiregulin receptor ERBB1 (35). Mice with global deletion of Areg (*Areg*^*-/-*^ mice), or with deletion of the *ErbB1* receptor expressed on cardiac myocytes (*ErbB1*^*loxP/loxP*^ Myosin Cre*+*) experienced increased susceptibility to myocardial ischemia and reperfusion injury. Together, those studies implicate myocyte-dependent HIF2A in coordinating the induction and signaling events of AREG through the ERBB1 receptor in cardioprotection (23, 35). Together with the current studies, those findings accentuate a cardioprotective role of HIF, by attenuating myocardial ischemic injury through myocyte-dependent HIF2A and myocardial reperfusion injury through myeloid-dependent HIF1A.

While the current studies concentrate on myocardial ischemia and reperfusion injury, previous studies had also implicated HIF in promoting ischemic preconditioning of the heart. Ischemic preconditioning involves short episode of ischemia that confers protection to a subsequent myocardial infarction (59). Several previous studies have shown functional roles of HIF1A in mediating the tissue-protective effects of ischemic preconditioning, including studies to address tissue-specific functions of HIF1A. Initial studies demonstrated that mice with partial deletion of *Hif1a* (*Hif1a*^*+/-*^ mice) experienced a complete loss of cardioprotection from ischemic preconditioning (26). Similarly, mice with siRNA in vivo silencing of Hif1a demonstrated a lack of cardio-protection by ischemic preconditioning, while pre-treatment with the HIF stabilizer DMOG preconditioned the myocardium (5). Subsequent studies to address tissue-specific functions of HIF1A in ischemic preconditioning implicated HIF1A activity in vascular endothelia (52). Other studies found that deletion of *Hif1a* in cardiomyocytes (*Hif1a*^*loxP/loxP*^ Myosin Cre*+*) also abolishes the cardio-protective effects of ischemic preconditioning (23). In contrast, mice with cardiomyocyte-specific deletion of *Hif2a* (*Hif2a*^*loxP/loxP*^ Myosin Cre*+*) were protected by ischemic preconditioning (23). Interestingly, more recent studies also implicate HIF1A in mediating the cardioprotective effects of remote ischemic preconditioning, where repeated episodes of remote ischemia to a limb can produce protection from myocardial or acute kidney injury (60). These studies indicate that during remote ischemic preconditioning, HIF1A is stabilized, and transcriptionally induces the HIF target gene L-10 with subsequent IL-10 signaling as a mechanism of cardioprotection (51). Taken together, these studies provide evidence from multiple studies that HIF1A – as opposed to HIF2A – mediates the cardioprotective effects of direct or remote ischemic preconditioning (17, 61).

The current studies implicate the HIF1A target netrin-1 in PMNs as critical target for mediating the cardioprotective effects of myeloid HIF1A. Netrin-1 is a neuronal guidance molecule that was initially described as a diffusible axon outgrowth promoting factor in nematodes (62). However, it became apparent that netrin-1 is much more ubiquitously expressed (63). In addition to its function as neuronal guidance molecule during brain development (47), it became apparent that netrin-1 can also function to modulate inflammatory endpoints or ischemia and reperfusion injury (64, 65). For example, netrin-1 released from the vagal nerve has been shown to attenuate excessive inflammation and promote the resolution of injury (65). Several previous studies have linked netrin-1 signaling in cardioprotection from ischemia and reperfusion injury, for example via interaction of netrin-1 with the classic netrin-1 receptor deleted in colorectal cancer DCC expressed on endothelial cells (66, 67). In line with the current findings, a recent study indicates that PMN-dependent netrin-1 can function to provide cardioprotection during in situ ischemia and reperfusion injury by enhancing extracellular adenosine signaling events through the Adora2b adenosine receptor (12, 64, 68–70). While extracellular adenosine signaling was initially described to induce transient slowing of the heart rate (71), many subsequent studies have sown functional roles for extracellular adenosine signaling in attenuating myocardial ischemia and reperfusion injury, including studies directly implicating myeloid Adora2b signaling (7, 38, 45). Taken together, those studies provide additional support for the concept that myeloid HIF1A provides cardioprotection via transcriptional induction of its target gene netrin-1, thereby leading to attenuated myocardial ischemia and reperfusion injury.

The current findings have important translational implications. For example, it is conceivable that patients experiencing myocardial ischemia and reperfusion injury could be treated with recombinant netrin-1 to activate the HIF1A-netrin-1 pathway for cardioprotection. Several other studies have shown that treatment with recombinant netrin-1 can function to dampen myocardial injury (12) or other inflammatory disease states (68). However, an alternative treatment approach would include the use of pharmacologic HIF stabilizers. Over the past decade, several pharmaceutical companies have developed pharmacologic HIF activators (17, 18, 72–74). These pharmacologic HIF activators function by inhibition of prolylhydroxylases, and thereby prevent the degradation of HIF1A via the proteasomal pathway (3). Pharmacologic HIF activators such as roxadustat or vadadustat have been successfully trialed for the treatment of renal anemia and phase 3 clinical trials and are available as oral medications (19–22). These orally available HIF activators could be used in patients experiencing myocardial ischemia and reperfusion injury. In addition, it is conceivable that these medications could be given to patients who are at high risk for experiencing myocardial injury, such as patients undergoing cardiac surgery. Treatment with an orally available HIF activator would likely cause HIF stabilization including in neutrophils, and thereby dampen myocardial ischemia and reperfusion injury (75). This highlights that HIF1A coordinates the transcriptional response in response to myocardial ischemia.

## Acknowledgements

All schematics were generated using BioRender.com

## Notes

**Funding Statement:** The present studies are supported by Deutsche Forschungsgemeinschaft (DFG; KO 3884/5-1) to M.K and National Institute of Health Grants R01HL154720, R01DK122796, R01DK109574, R01HL133900 and Department of Defense Grant W81XWH2110032 to HKE

**Conflicts of Interest:** The authors declare no competing interests.

### Competing Interest Statement

The authors have declared no competing interest.

